# Mesh convergence depends on the element formulation of finite element brain models

**DOI:** 10.64898/2026.07.30.741810

**Authors:** Azilis Even, Zhou Zhou, Svein Kleiven

## Abstract

Finite element (FE) head models are virtual tools to study brain biomechanics and their predictions must be numerically convergent. Previous convergence studies focused on the influence of mesh size, but the potential effect of element formulation on model convergence was often ignored. To address this, one original model with brain mesh size as 6.4 ± 1.9 mm was modified to generate three derivatives with the same mesh topology but different element sizes, i.e., a coarse model (mesh size: 12.2 ± 3.9 mm), a medium model (mesh size: 3.2 ± 1.0 mm), and a fine model (mesh size: 1.6 ± 0.5 mm). Three commonly used element formulations, i.e., reduced integration, selectively reduced (S/R) integration, and full integration, were implemented to the brain elements. These models were subjected to rotational loadings along the axial, coronal, and sagittal axes, respectively. The maximum relative displacement at representative sites and 95^th^ percentile maximum principal strain at the whole brain level were used to evaluate mesh convergency. The results showed that the S/R integration yielded a 5% difference between the original and medium meshes, while the reduced and full integration revealed a difference over 5% even between the medium and fine meshes. This study verified that the mesh convergence of FE brain models is affected by the choice of element formulation and the S/R integration contributes to the fastest convergence behavior than the reduced and full integrations. It provided practical information on how to develop numerically convergent and computationally efficient FE brain models.

**Highlights:**

- This study verifies that the choice of element formulation affects the mesh convergence behavior of finite element brain models
- This study finds the selectively reduced integration yields the fastest convergence behavior than the reduced and full integration
- This study provides practical guidance on the choice of mesh density and element formulation on how to develop numerically convergent and computationally efficient finite element brain models.

## Introduction

Traumatic brain injury (TBI) is a major public health issue associated with high rates of incidence, mortality, and morbidity. Globally, an estimated 69 million people sustain a TBI each year [1], resulting in an annual economic cost of approximately 400 billion dollars [2]. Nationally, the overall TBI incidence was 852 per 100,000 person-years in New Zealand [3], and 343 and 285 per 100,000 person- years in Sweden for men and women respectively [4], with falls and traffic accidents being the dominant causes of TBI in both countries. In the specific case of traffic accidents, TBI was shown to be a major source of injury for vulnerable road users, with pedestrians suffering from severe TBI more often than cyclists and powered two-wheeler riders [5]. Understanding the complex mechanisms of TBI is crucial to addressing the significant socioeconomic burden it represents.

Finite element (FE) models are computational surrogates for virtual analysis, simulation, and prediction of a real-world object or process, such as estimating internal brain tissue responses secondary to external head impacts. To ensure reliable assessment, an FE head model must have sufficient representation of various intracranial components in terms of geometry [6], material [7], and interface [8] and needs to be appropriately validated against experimental measurements pertaining to its intended application [9–11]. One important modelling variable is the mesh density, which determines the characteristic element size and, consequently, the number of elements in the FE head model. An appropriate mesh density should achieve convergent response predictions - such that further mesh refinement does not meaningfully alter the predicted response (so-called mesh convergence) - while maintaining acceptable computational efficiency [12, 13]. Existing convergence studies for 3D head models evaluated the effect of element size on the brain response using criteria such as pressure [14, 15], brain displacements [12, 14–16], and more recently brain strains [13, 17, 18], although the pressure has now been understood to be less relevant to injury [12]. Despite these previous investigations, it appears no consensus has yet been reached regarding the mesh density required for convergent brain response predictions, indicating that numerical factors beyond mesh density may also influence the convergence characteristics of FE head models.

Element formulation is another key modelling parameter. By definition, it determines both the element shape function (i.e. how the displacement is interpolated over the elements) and the integration scheme (i.e. which integration points are used to evaluate stress and strain over the elements) [19]. It is known that the element formulation choice can influence the model stability, the computational cost of impact simulations and the model-estimated brain response [16, 17]. Current hexahedron-based FE head models vary in terms of element formulation, with available options including reduced, selectively reduced (S/R), and full integration 8-node elements (see the method section for details) [19, 20]. When expanding to the context of mesh convergence, existing studies often implemented a fixed element formulation as defined in a specific FE solver, including reduced integration [17] and S/R integration [12, 14] in LS-Dyna, or C3D8I element formulation (incompatible modes 8-node brick element) in Abaqus [13]. Consequently, how element formulation interacts with mesh size to influence the predicted brain response and the associated assessment of mesh convergence remains poorly understood.

This study aimed to evaluate the influence of element formulation on the mesh convergence behavior of FE brain models. Three element formulations were implemented to four models with identical geometric profiles and mesh topologies but different average element sizes. These twelve models were used to evaluate the convergence behavior in brain displacements and strain during head impacts. By comparing the model-estimated brain displacement and strain, the hypothesis that the mesh convergence depends on the element formulation of FE brain models was evaluated.

## Methods

### Brain meshes and element formulations

The KTH head model [21] served as the baseline for this study. Its brain has 4212 elements and is shown in Figure 1 (labelled as original mesh). To investigate the mesh convergence, three new brain meshes were derived from this original model by either merging or splitting its elements. The four resulting brain models had the same mesh topology but different element sizes (Figure 1). They included a coarse model with average mesh size 12.2 ± 3.9 mm, the original KTH head model with mesh size 6.4 ± 1.9 mm, a medium model with mesh size 3.2 ± 0.97 mm and a fine model with mesh size 1.6 ± 0.48 mm. The mesh quality of the meshes included in this study was evaluated using several criteria [13, 21], with the element quality being comparable among all four meshes (Table 1).

**Figure 1.**
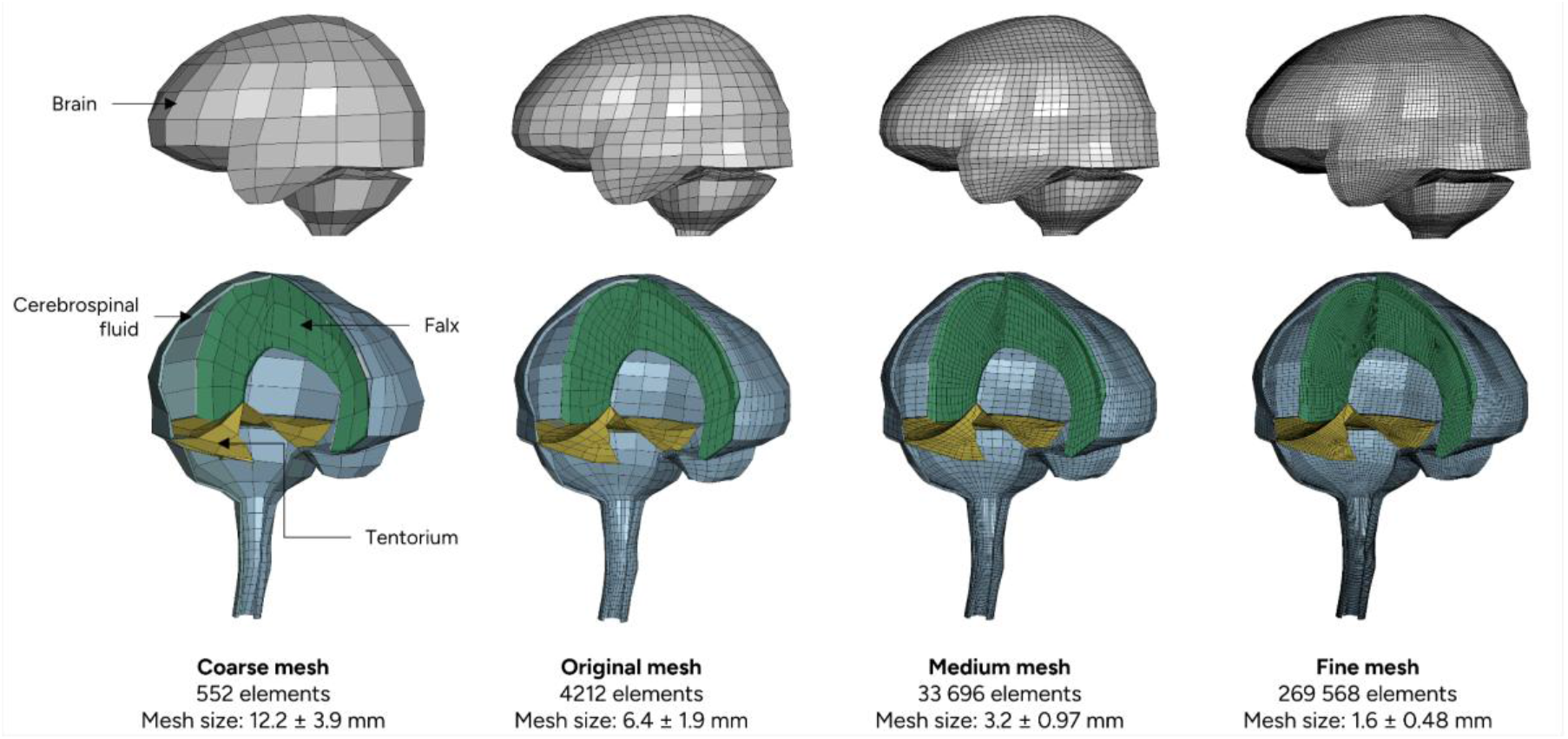
Four versions of the finite element brain models with different mesh densities. For clarity, a portion of cerebrospinal fluid elements were masked in the low-row subfigures to better visualize the falx and tentorium.

**Table 1.**
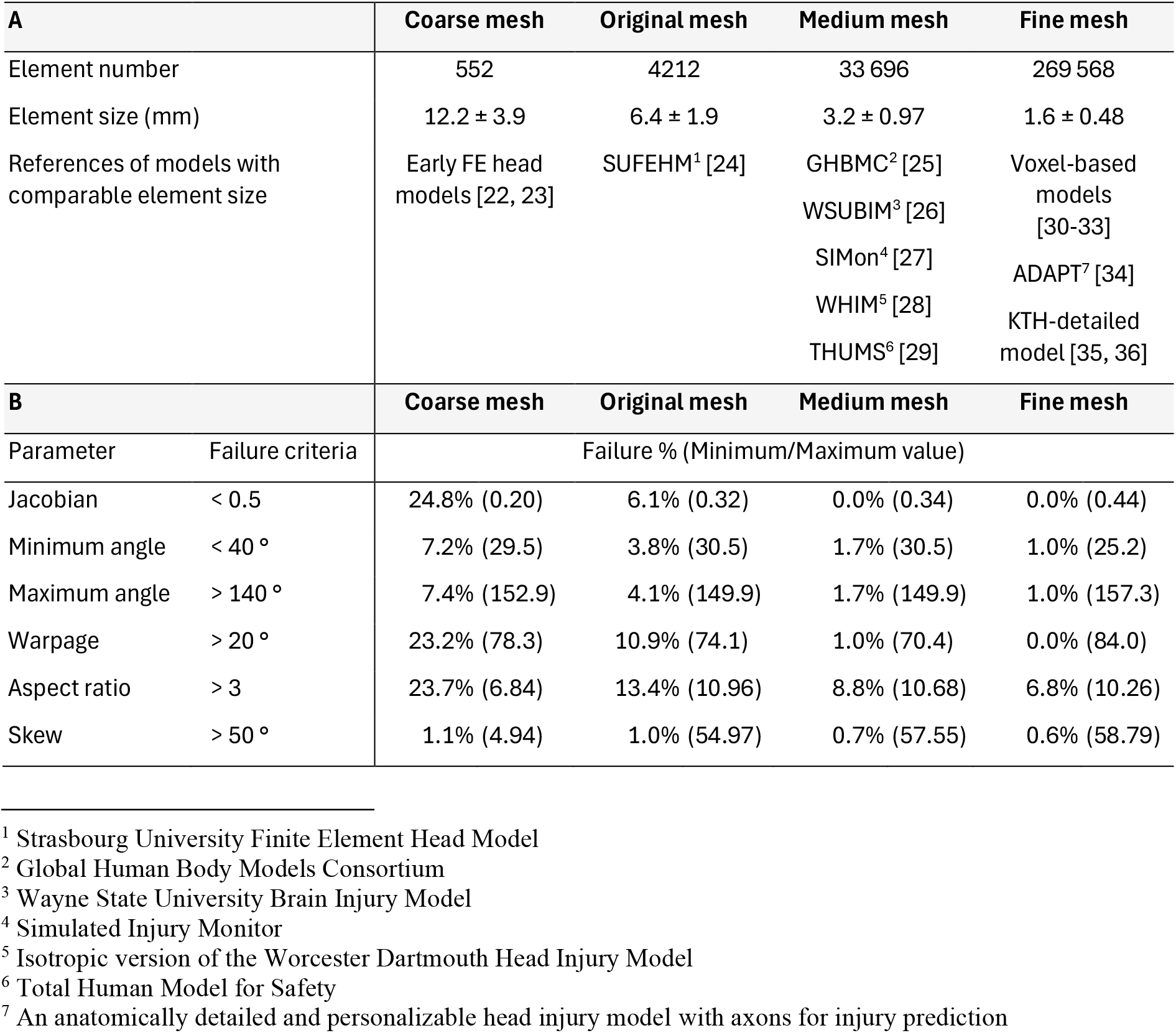
Summary of brain element number and size (A) and mesh quality (B) of the four brain meshes.

Various element formulations were applied to the four meshes described above to investigate the potential influence of element formulation on model convergence. Specifically, the reduced integration elements, S/R integration elements, and full integration quadratic elements with nodal rotations were considered, motivated by their use within the brain modelling community. The reduced integration elements, alternatively termed as constant stress solid elements, use only one point of integration at the centroid of the element. This formulation is the least computationally expensive and is used in most available FE head models, including THUMS [29], SIMon [27], GHBMC [25] and the isotropic WHIM [28]. It is, however, under-integrated, which has the potential to create unwanted zero-energy modes (also referred to as hourglass modes) that need to be controlled to limit the risk of numerical instabilities [37, 38]. Full integration elements are prone to both volumetric locking problems when using nearly incompressible materials and shear locking problems when using poor aspect ratio elements [37]. No commonly available hexahedra-based FE head models implement this formulation. The S/R formulation positions itself as an in-between option and is used in the KTH head Model [21] and other counterparts [39]. It splits the strains into deviatoric and volumetric strains and uses a full integration scheme for the deviatoric strains and a reduced integration scheme for the volumetric strains. Both the full and S/R integration have no hourglass issue. In this study, LS-Dyna (version R15.0, Ansys Inc.) was used to apply these three integrations to the brain meshes. Specifically, the S/R integration was implemented using the “ELFORM = 2” option and the full integration quadratic elements with nodal rotations using “ELFORM = 3” [37, 40]. The reduced integration was implemented using the “ELFORM = 1” option combined with a Flanagan-Belytschko viscous form of hourglass control with exact volume integration [38], similar to the choice of the GHBMC and SIMon models [17, 25].

For the material of resultant twelve models (i.e. three element formulations implemented on four brain meshes), the brain was all modelled as a homogeneous structure with nonlinear behavior using a second-order Ogden-based hyperelastic constitutive law with six Prony-series terms to account for its rate dependency. The cerebrospinal fluid was modelled as a nearly incompressible elastic fluid [21, 34, 39]. Other structures were modelled according to Table 2. It should be noted that two additional brain material parameters were implemented to the twelve models with the purpose of evaluating the potential influence of material properties on mesh convergence. The corresponding results are presented in Appendix A and demonstrate that the main conclusions remain unchanged.

**Table 2.** Material properties of each component in FE head models. Note that *μ_i_* and *α_i_* are Ogden parameters, *G_i_* represents the 6 shear relaxation moduli, *β_i_* are the 6 decay constants.

| Material | Description |  |
| --- | --- | --- |
| Brain | Ogden parameters | $\mu_1=53.8$ Pa, $\alpha_1=10.1$,<br>$\mu_2=-120.4$ Pa, $\alpha_2=-12.9$ |
| | Shear relaxation modules | $G_1=0.31$ MPa, $G_2=78$ kPa,<br>$G_3=6.2$ kPa, $G_4=8.0$ kPa,<br>$G_5=1.0$ kPa, $G_6=3.0$ kPa |
| | Decay constants | $\beta_1 \cdots \beta_6 = 10^6 \cdots 10^1$ (1/s) |
| Cerebrospinal fluid | Elastic fluid with the bulk modulus as 2.1 GPa |  |
| Falx & tentorium | Simplified rubber with experimentally derived stress-strain curve [41] |  |
| Pia mater | Simplified rubber with experimentally derived stress-strain curve [42] |  |
| Skull & dura mater | Rigid |  |

### Impact simulations

Whole-head impact simulations were performed using the twelve models to estimate brain responses. Three rotational velocity loading profiles, all of which were nearly half-sine shaped with a peak of approximately 40 rad/s and a duration of approximately 30 ms, were applied to the twelve models along three anatomical axes (i.e., axial, coronal, and sagittal), respectively. These loading profiles were obtained from the controlled laboratory cadaveric experiments reported by Alshareef, et al. [43], specifically from specimen 846M, in which brain displacement was measured under nearly pure rotational loading conditions. In each simulation, the loading profile was imposed on a node at the center of gravity that was attached to the rigid skull. All simulations were solved using LS-Dyna MPP (massively parallel processing) version R15.0.

### Evaluation of the convergence behavior

The convergence behavior was evaluated by comparing the maximum resultant displacement relative to the skull for the six representative nodes inside the brain for all models (Figure 2). These nodes were chosen based on their locations in the coarse model less than 5 mm away from those of the sonomicrometry crystals from Alshareef, et al. [43]. Note that the nomenclature of those nodes in Figure 2 was the same as that in the original experimental study.

**Figure 2.**
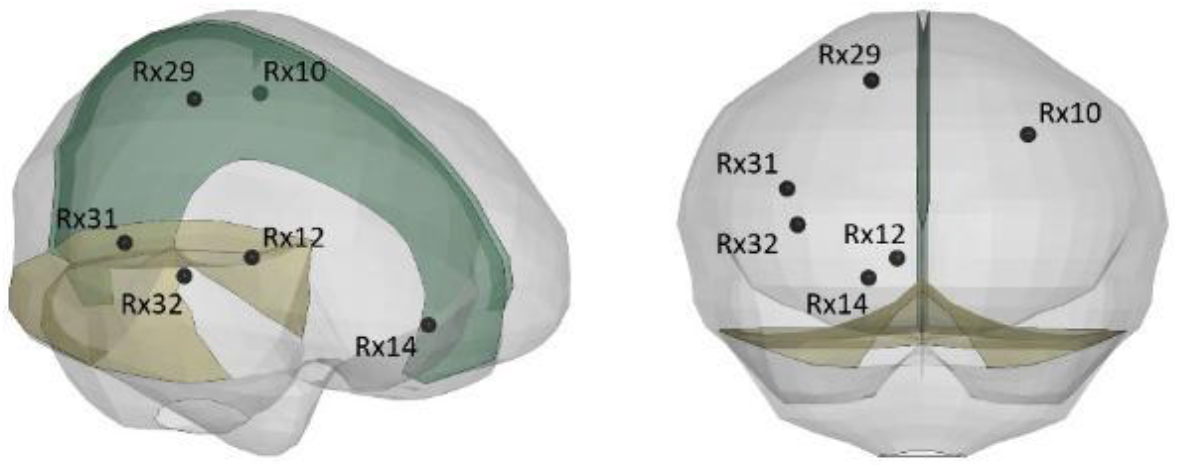
Isometric and frontal views of the locations of 6 representative nodes used to derive brain displacement responses. Note that, to improve visualization of the node locations, the pia mater (light gray), falx (light green), and tentorium (light yellow) are rendered with transparency.

To evaluate the effect of element formulation on strain-based convergence, the 95^th^ percentile maximum principal strain (MPS) was extracted. For each element of the brain mesh, the MPS was defined as the peak first principal Green-Lagrange strain over the simulation duration, and the 95^th^ percentile over the whole brain was then considered. The top 5% of peak element values were excluded on the assumption that they could suffer from numerical instabilities [39, 44]. The absolute change in strain for each step of mesh refinement was evaluated. For each simulation, the volume fraction of brain elements over a varying MPS threshold was examined to get a detailed overview of the strain distribution across the brain depending on mesh density and element formulation [39, 45, 46].

## Results

### Evaluation of model convergence based on brain displacements

Maximum relative displacements of six representative nodes were used to evaluate the model convergence (Figure 3). Across all three loading conditions, the models employing the S/R integration consistently predicted larger maximum relative displacements than those using the reduced integration, while the full integration formulation generally predicted the smallest values. For the models with fine meshes, all three element formulations were shown to approach similar displacement results. The complete time-history curves of the resultant nodal displacement for each model are available in Appendix B.

**Figure 3.**
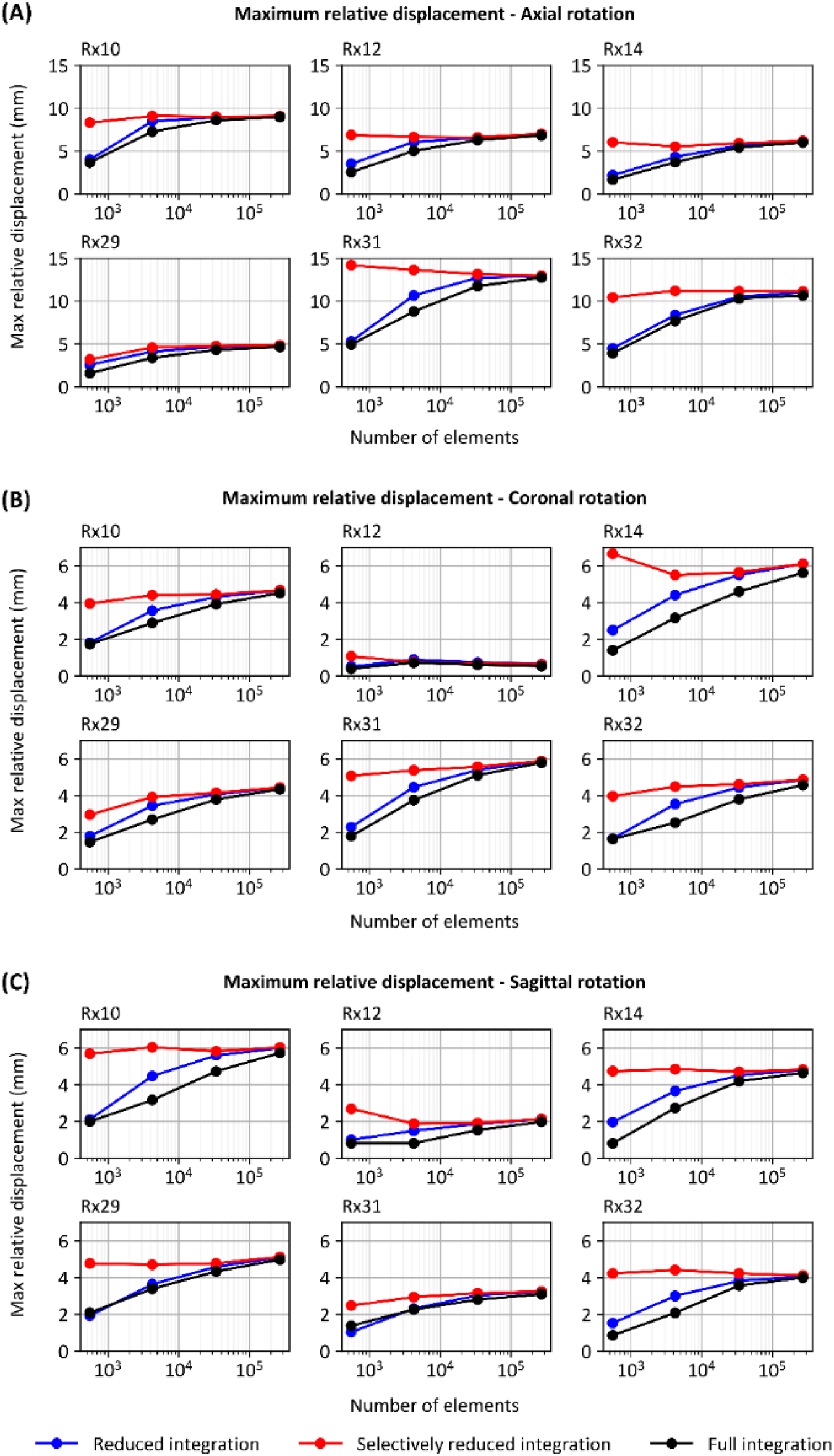
Maximum relative displacement of the six representative nodes for axial rotation (A), coronal rotation (B), and sagittal rotation (C). Note that for each dotted curve, the four dots from the left to the right represent the model with coarse mesh, original mesh, medium mesh, and fine mesh, respectively.

The S/R formulation exhibited the fastest convergence, regardless of the loading conditions. For the axial loading simulations implementing the S/R integration (Figure 3A), five out of six representative nodes showed a change in maximum relative displacement below 5% when refining from the original to medium mesh. With the reduced and full integration, changes in nodal displacements ranged from 5 to 50% when refining from the original to medium mesh. For the coronal loading simulations using the S/R element formulation (Figure 3B), four out of six representative nodes showed a change in maximum relative displacement below 5% when refining from the original to medium mesh. In contrast, with the reduced and full integration, all changes in nodal displacements were over 20% when refining from the original to medium mesh, and over 5% when refining from the medium to fine mesh. For the sagittal loading simulations, only one node (i.e., Rx31) failed to meet the 5% threshold when refining from the original to medium mesh and using the S/R integration. The reduced and full integration showed slower convergence behavior, with changes in nodal displacement remaining over 5% when refining from the medium to fine mesh. For reference, all the calculated changes in maximum nodal displacements are available in Appendix C.

### Evaluation of model convergence based on brain strain

The combined influence of mesh density and element formulation on brain strain peaks is described in Figure 4. For all three loadings, when implementing the S/R element formulation, the change in 95^th^ percentile MPS was below 25% when refining from the coarse to original mesh, and below 5% when refining from the original to medium mesh. With the reduced integration formulation, the change in strain was below 10% for all three loading cases when refining from the medium to fine mesh but was significantly higher for all other refinement steps (e.g., 21.9% between the original and medium meshes under coronal rotation). With full integration elements, the change in strain was over 5% for all refinement steps and all three loading cases. For reference, all the calculated changes in 95^th^ percentile MPS values are available in Appendix D. Furthermore, the brain strain values of other percentiles are presented in Appendix E, further reinforcing that the S/R integration contributed to the fastest convergence than the reduced and full integration.

**Figure 4.**
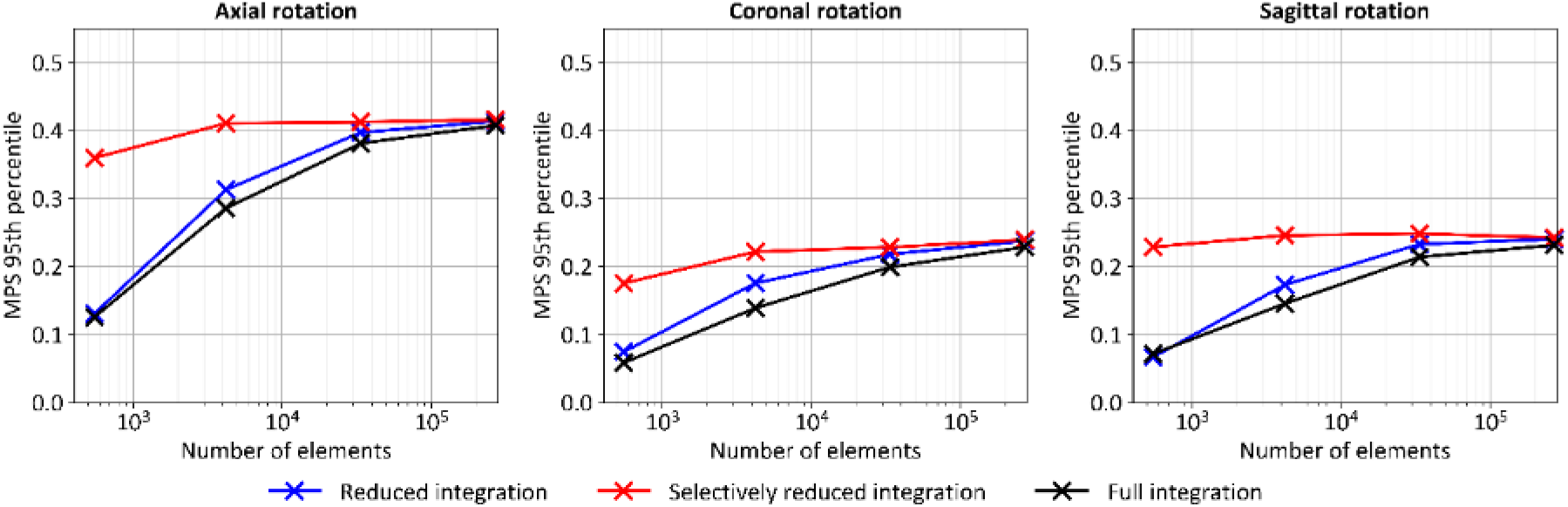
Comparison of the 95th percentile maximum principal strain (MPS) of the brain under axial, coronal, and sagittal rotations, illustrating the influence of mesh density and element formulation on the predicted peak brain strain.

We next examined the influence of mesh density and element formulation on brain strain distribution (Figure 5). For all three loading cases, when using the S/R integration, the strain contours for the original mesh and the finer meshes were visually similar (Figure 5A, C, E). When switching to the reduced and full integration formulations, the area of high strain (characterized by a red color in the contour) estimated by the models with the original and medium meshes were different. We further quantitatively calculated the cumulative volume fraction results above varying strain levels (Figure 5B, D, F). For the S/R formulation, the cumulative volume fraction-strain curves obtained with the original mesh were nearly superimposed on those of the medium and fine meshes in all three loading cases. In contrast, the reduced and full integration formulations exhibited systematic differences, with the original mesh consistently predicting smaller brain volume fractions for a given strain threshold than the medium and fine meshes.

**Figure 5.**
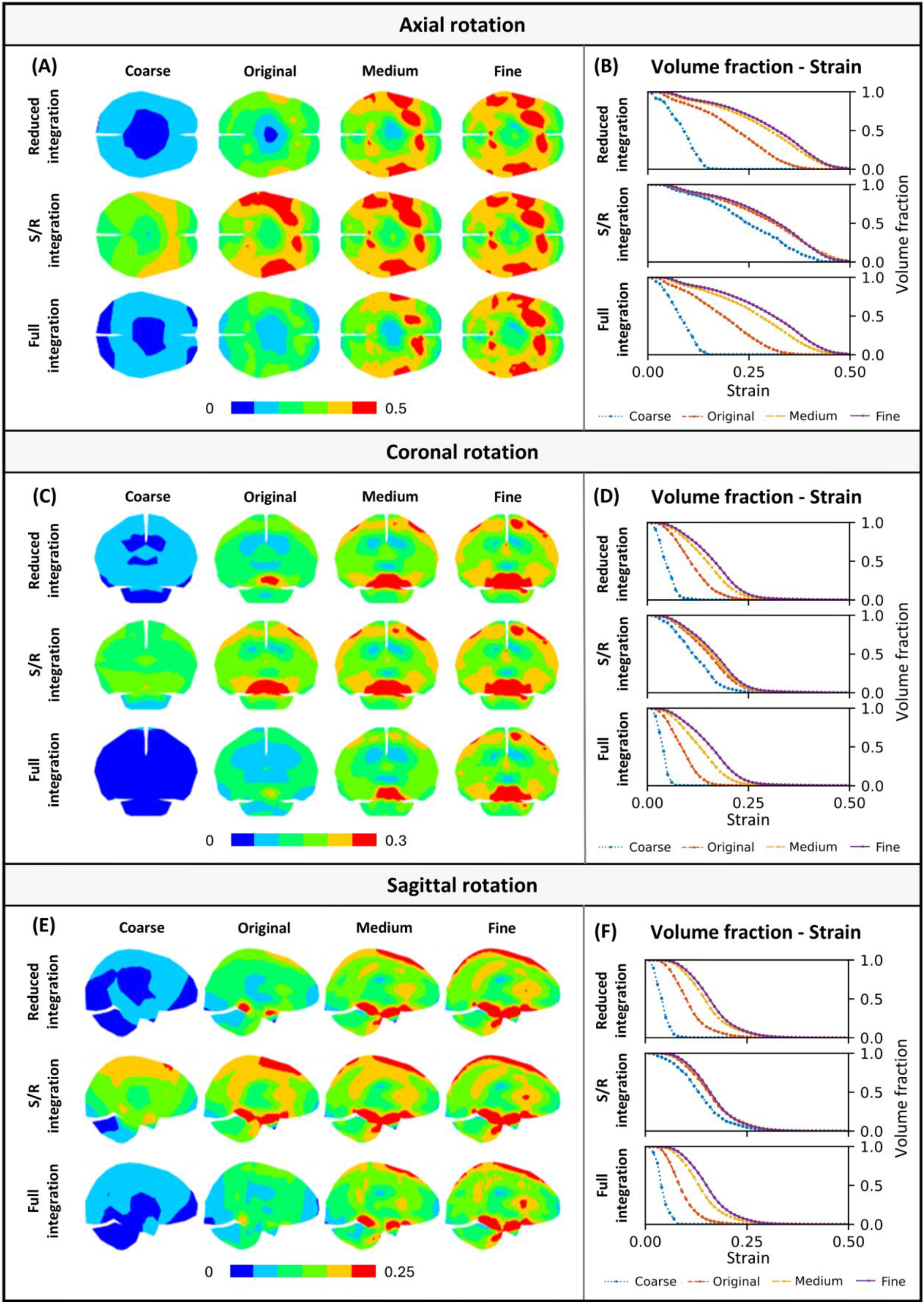
Element-wise peak strain contour and volume fraction of the brain elements with the strain over varying thresholds for the axial (a-b), coronal (c-d) and sagittal (e-f) loadings. For each loading, the strain contour is shown on the left, and the volume fraction-strain curves are on the right.

### Computational time and hourglass energy ratio

We employed a representative simulation (i.e., axial rotation with the simulation time of 30 ms) to illustrate the influence of mesh size and element formulation on the computational cost. As shown in Table 3A, the required computational time increased with the mesh density and was affected by the element formulation. For a given mesh, reduced integration elements exhibited the lowest computational cost, followed by S/R integration elements, while full integration elements were associated with the highest computational cost.

**Table 3.**
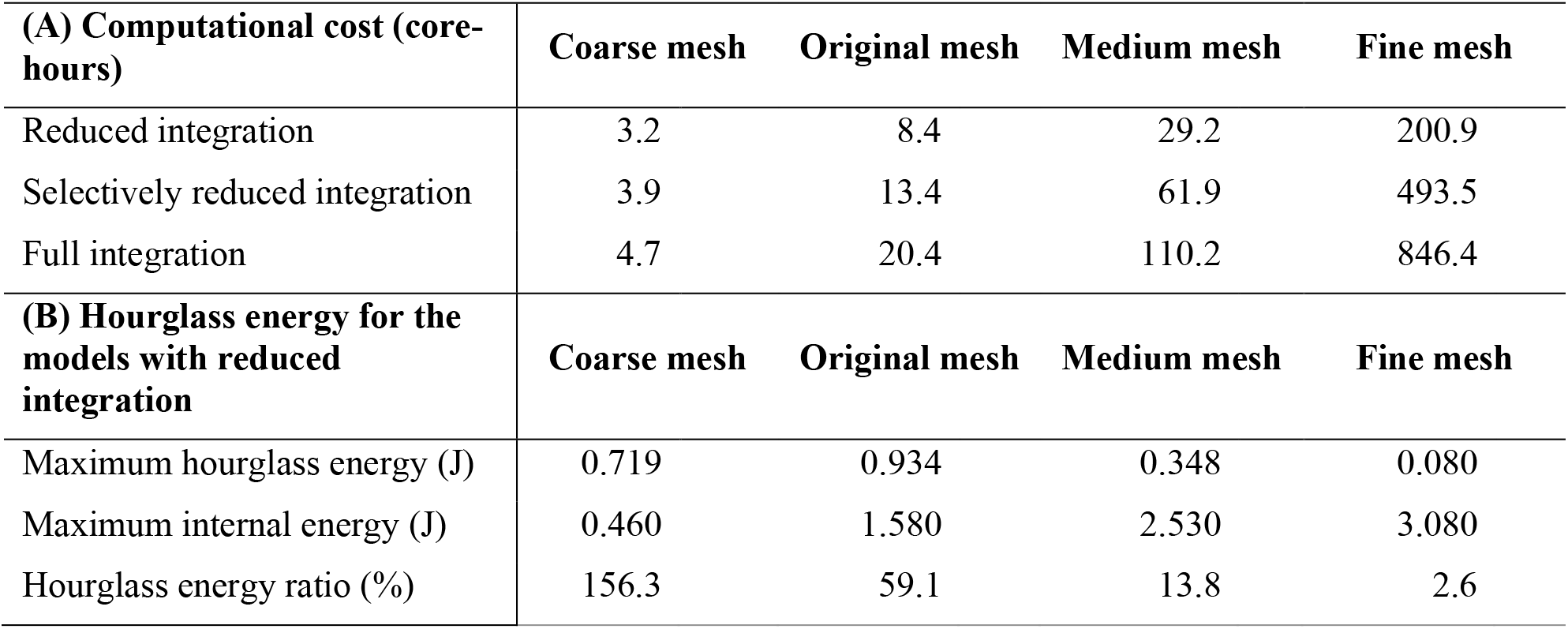
(A) Computational cost (in core-hours) of 12 models (covering four mesh densities and three element formulations) for the simulation of axial rotation. (B) The hourglass energy ratio (calculated as the ratio of the maximum hourglass energy and the maximum internal energy) of the four models with reduced integration for the simulation of axial rotation.

| **(A) Computational cost (core-hours)** | **Coarse mesh** | **Original mesh** | **Medium mesh** | **Fine mesh** |
| --- | --- | --- | --- | --- |
| Reduced integration | 3.2 | 8.4 | 29.2 | 200.9 |
| Selectively reduced integration | 3.9 | 13.4 | 61.9 | 493.5 |
| Full integration | 4.7 | 20.4 | 110.2 | 846.4 |
| **(B) Hourglass energy for the models with reduced integration** | **Coarse mesh** | **Original mesh** | **Medium mesh** | **Fine mesh** |
| Maximum hourglass energy (J) | 0.719 | 0.934 | 0.348 | 0.080 |
| Maximum internal energy (J) | 0.460 | 1.580 | 2.530 | 3.080 |
| Hourglass energy ratio (%) | 156.3 | 59.1 | 13.8 | 2.6 |

To evaluate the stability of models implementing reduced integration, the hourglass energy ratio was calculated as the ratio of the maximum hourglass energy and the maximum internal energy (Table 3B). The hourglass energy ratio was also affected by the mesh density and decreased with the average element size. The hourglass energy ratio was over 10% when using the coarse, original and medium meshes, and under 5% when using the fine mesh.

## Discussion

This study implemented three element formulations (i.e., reduced integration, S/R integration, and full integration) to four FE head models with varying mesh densities (12.2 ± 3.9 mm ∼ 1.6 ± 0.48 mm) to evaluate how the element formulation affected the model convergence behavior. Our results showed that the S/R integration yielded the fastest convergence results both in terms of brain nodal displacements and brain strain, followed by the reduced integration and full integration. These findings verified the hypothesis that the mesh convergence of the FE brain model was affected by the element formulation. Our work provided practical information on the choice of mesh density and element formulation on how to develop numerically convergent and computationally efficient FE brain models.

The present study could be regarded as a significant extension of two previous works and provided novel insights on the combined influence of mesh density and element formulation. In one early study, Giudice, et al. [17] evaluated the influence of several numerical modelling choices, including mesh size and element formulation, through parametric studies. While their work demonstrated that both parameters substantially affected predicted brain responses, the element formulation study was limited to the reduced and full integration because simulations using the S/R integration terminated with error. The present study overcomes this by successfully implementing all three formulations within the same framework, enabling the new finding that the S/R formulation exhibited the most favorable mesh convergence characteristics. In contrast, Zhao and Ji [13] similarly conducted a mesh convergence study of the WHIM model in Abaqus using the C3D8I element formulation (incompatible modes 8-node brick element), reporting that an average element size of 1.8 mm was needed to achieve converged responses. However, we are unable to confidently identify the element formulation in LS-Dyna that is directly equivalent to the C3D8I element in Abaqus, and hence further comparison was not performed.

The presented results collectively investigated the influence of element formulation and mesh size on the brain response, and our findings are generally consistent with previous studies focusing on a single parameter (i.e., exclusively focused on either element formulation or mesh size). For the influence of element formulation, our work showed that, for a given mesh, the full integration generally led to the lowest peak displacements and strains, followed by the reduced integration, while the S/R integration revealed the highest brain deformation results (Figures 3-5). This was in agreement with the finding by Giudice, et al. [17], who found that switching the element formulation in the SIMon head (element size: ∼ 3.2 mm) from the reduced integration to the full integration lowered nodal displacements and MPS. For the influence of mesh size, early works were primarily conducted with the reduced integration formulation and reported that coarser meshes had a general tendency to underestimate brain displacement [15, 16] and brain strain [17]. Such a tendency was also noted in our simulation results using the reduced integration, of which the differences between the original and medium meshes were up to 30.8% in brain displacement (Appendix C) and 30.9% in brain strain peaks (Appendix D).

Interestingly, our results suggested strain-based metrics were more sensitive to mesh size than displacement-based metrics. For instance, coarse-to-original differences in nodal displacement ranged from approximately 50-140% for the reduced and full integration formulations, and from 3.5-45% for the S/R formulation. In comparison, the corresponding differences in strain ranged from approximately 120-170% for the reduced and full integration formulations, and from 7.5-22% for the S/R formulation (Appendices C-D). This slower convergence of strain compared with displacement was expected because strain is obtained from the spatial derivatives of the displacement field, making it inherently more sensitive to spatial discretization [47]. Given the prevalence of strain-based metrics for TBI investigations, we recommended future work to perform mesh convergence study with both strain and displacement examined.

Our work yielded important practical implications for computational brain modelling. From the perspective of mesh convergence, the S/R integration was a preferred element formulation choice for the FE model with an element size of around 6.4 mm (i.e., the original mesh in the current study, the same as the original KTH head model [14]). This was supported by our finding that the differences in terms of brain strain and nodal displacement estimated by the model with original mesh to its heavily discretized counterparts was generally less than 5% (Figures 3-5). These findings hence provided *a posteriori* support for the choice of the S/R integration in the original KTH head model from a mesh convergence perspective. The S/R integration was indeed developed to relax the excessive constraints causing volumetric locking when using fully integrated low-order elements by applying reduced integration to the volumetric strains only [48, 49]. In contrast to the reduced formulation, this approach is less susceptible to hourglass modes since it avoids the rank deficiency caused by one-point integration [38]. Shear locking remains a potential issue in structures using poor aspect ratio elements [20, 37]. For FE models with a relatively higher mesh resolution (e.g., the fine mesh with an averaged element size of 1.6 mm herein), the reduced integration represents a computationally efficient element formulation without compromising mesh convergence or introducing significant hourglass effects. This is supported by the finding that, for the fine mesh, the reduced and S/R integration produced virtually identical brain responses, while the computational cost of the former was less than half of the latter (Table 3A). Equally importantly, the hourglass energy ratio of the fine mesh using the reduced integration was approximately 2.6% (Table 3B), well below the 10% threshold previously mentioned by Giudice, et al. [17]. Indeed, several brain FE models with an element size comparable to the fine mesh or even finer used the reduced integration as the element formulation choice [31, 35, 50]. In contrast, the full integration did not appear to be a recommended choice for computational brain modelling, as it has slower convergence behavior and was computationally more expensive. This point has been discussed extensively elsewhere [17].

We focused on brain displacement and brain strain as the main model responses to evaluate mesh convergence. Future studies could extend this assessment to other injury metrics, such as the axonal fiber strain that has been recently used as injury metrics [18, 34, 45, 46, 51]. Garimella and Kraft [18] took an initial step in this direction by performing a mesh convergence study with an embedded truss representation of white matter fiber bundles to establish the mesh density required for converged axonal strain predictions.

## Conclusion

The current study implemented three element formulations to four brain models with the same mesh topology but varying element sizes to investigate the model’s mesh convergence behavior. By comparing the numerically predicted brain-skull relative displacement and brain strain, it was verified that the element formulation choice affects the mesh convergence outcome. The selectively reduced integration element formulation provides the fastest mesh convergence behavior than the reduced integration and full integration. Our work provided practical implications on the choice of mesh density and element formulation for the development of numerically convergent and computationally efficient FE brain models.

## Acknowledgements

This work was supported by FFI (Strategic Vehicle Research and Innovation), project number 2024- 03635, funded by Vinnova, the Swedish Transport Administration, the Swedish Energy Agency, and the industrial partners. We acknowledge Professor Matthew Panzer from the University of Virginia for providing the experimental data that was used in the current study. We also thank Prof. Xiaogai Li from KTH and Dr. Shiyang Meng from Autoliv for commenting on the early version of this manuscript. The content of this article is solely the responsibility of the authors and does not necessarily represent the official views of funding agencies. The computational simulations were enabled by resources provided by the National Academic Infrastructure for Supercomputing in Sweden (NAISS) at the Center for High Performance Computing (PDC) partially funded by the Swedish Research Council through grant agreement no. 2022-06725.

## Appendix A. Influence of the brain material properties on the model convergence behavior

To evaluate whether the brain stiffness affected the mesh convergence outcome, two additional material parameter sets were obtained by halving and doubling the corresponding values used in Table 2. The resultant two new sets of brain material properties are shown in Table A1 (respectively referred to as softer brain and stiffer brain) and were implemented to the twelve models (i.e., three element formulations implemented on four brain meshes) to estimate the brain response secondary to the loading conditions detailed in “Impact simulation” section. The predicted volume fraction of brain elements over certain strain thresholds are plotted in Figure A1 for the “softer brain” and in Figure A2 for the “stiffer brain”. It was clearly shown that, independent of the brain stiffness, the selectively reduced integration led to the fastest converged result in terms of volume fraction of brain strain across all three rotational directions than the reduced and full integration.

**Table A1.**
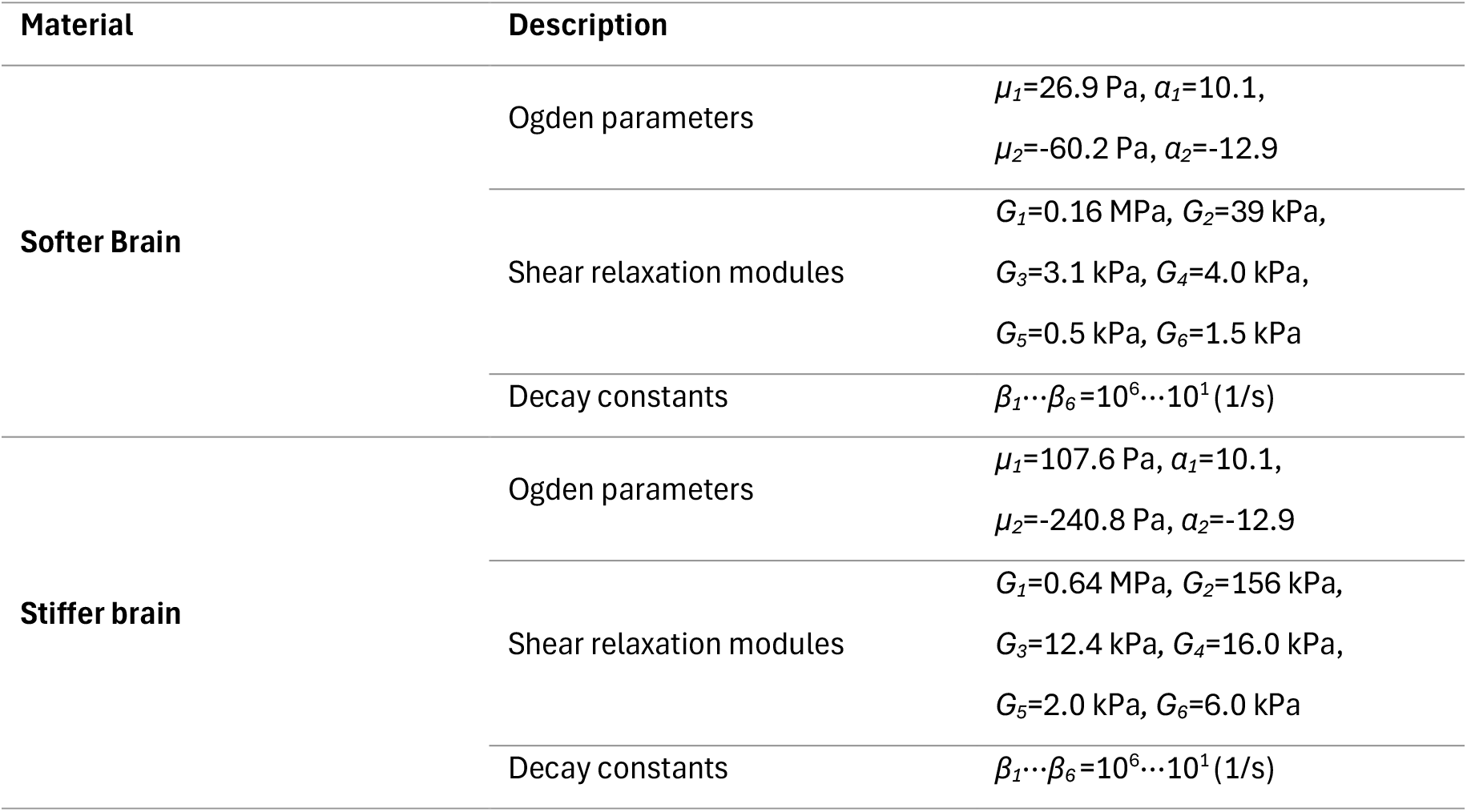
Material properties of the brain with soft and stiffer properties. Note that *μ_i_* and *α_i_* are Ogden parameters, *G_i_* represents the 6 shear relaxation moduli, *β_i_* are the 6 decay constants.

**Figure A1.**
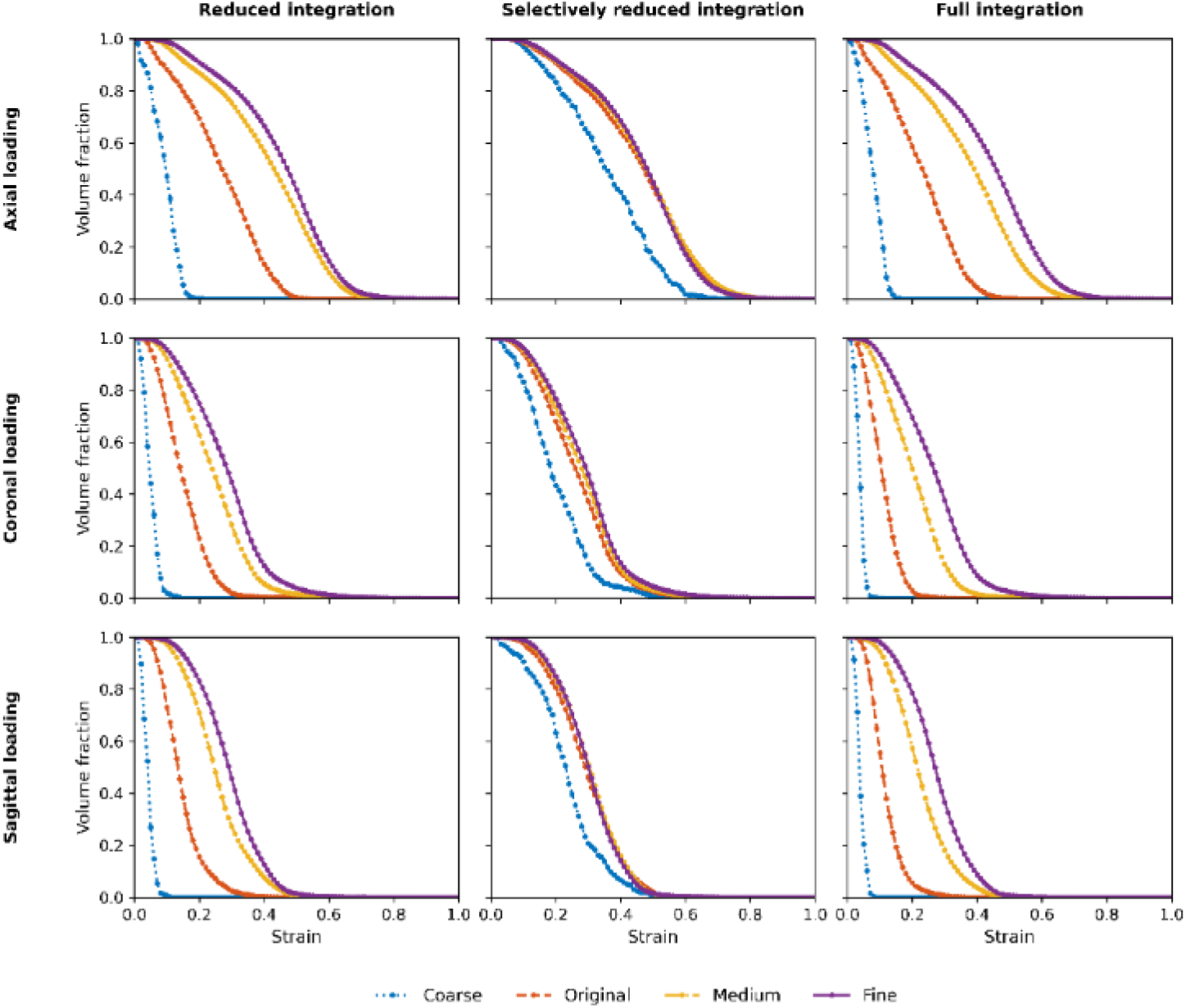
Volume fraction of the elements with the strain over varying thresholds for softer brain parameters.

**Figure A2.**
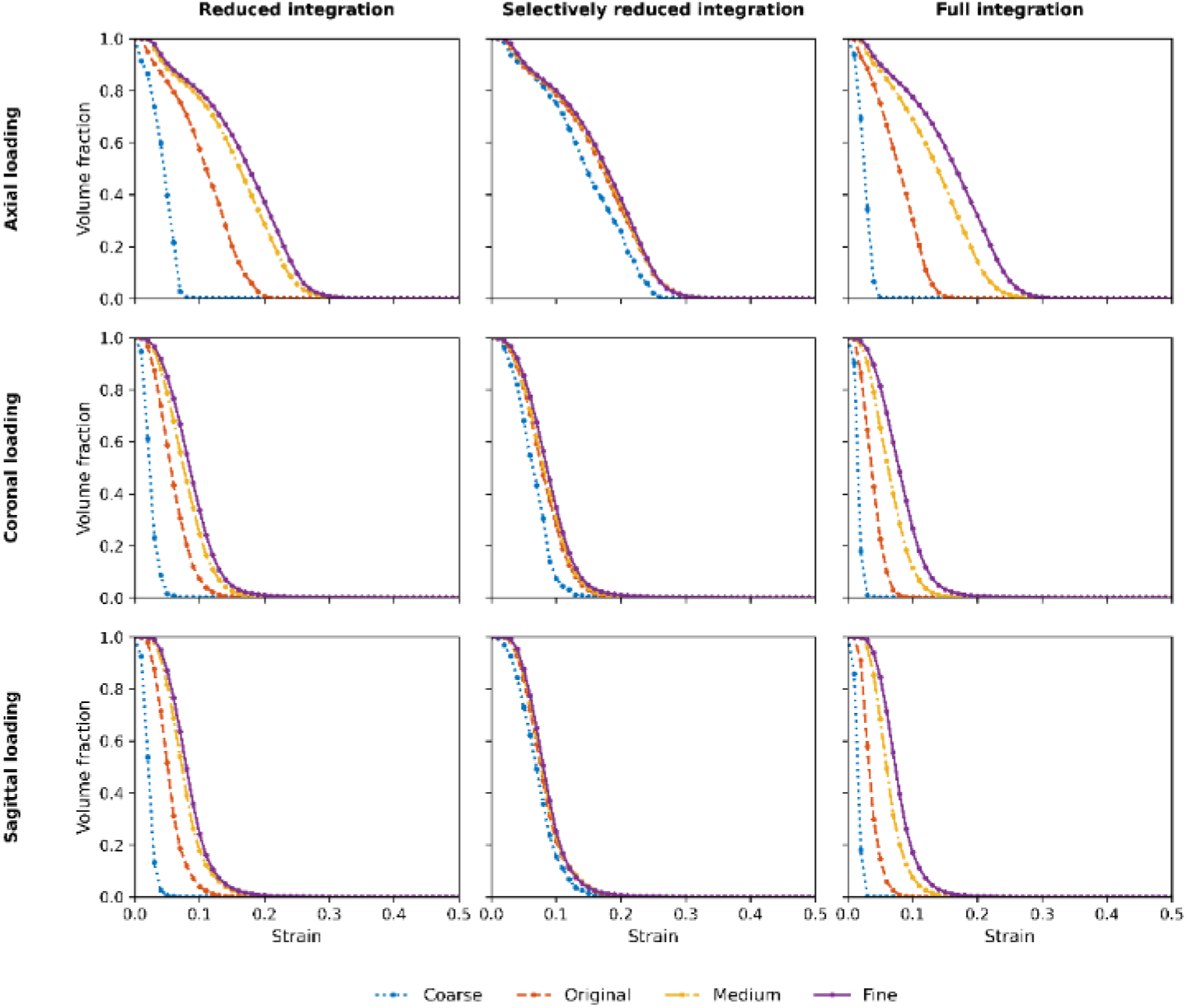
Volume fraction of the elements with the strain over varying thresholds for stiffer brain parameters.

## Appendix B. Nodal displacement time-history curves

**Figure B1.**
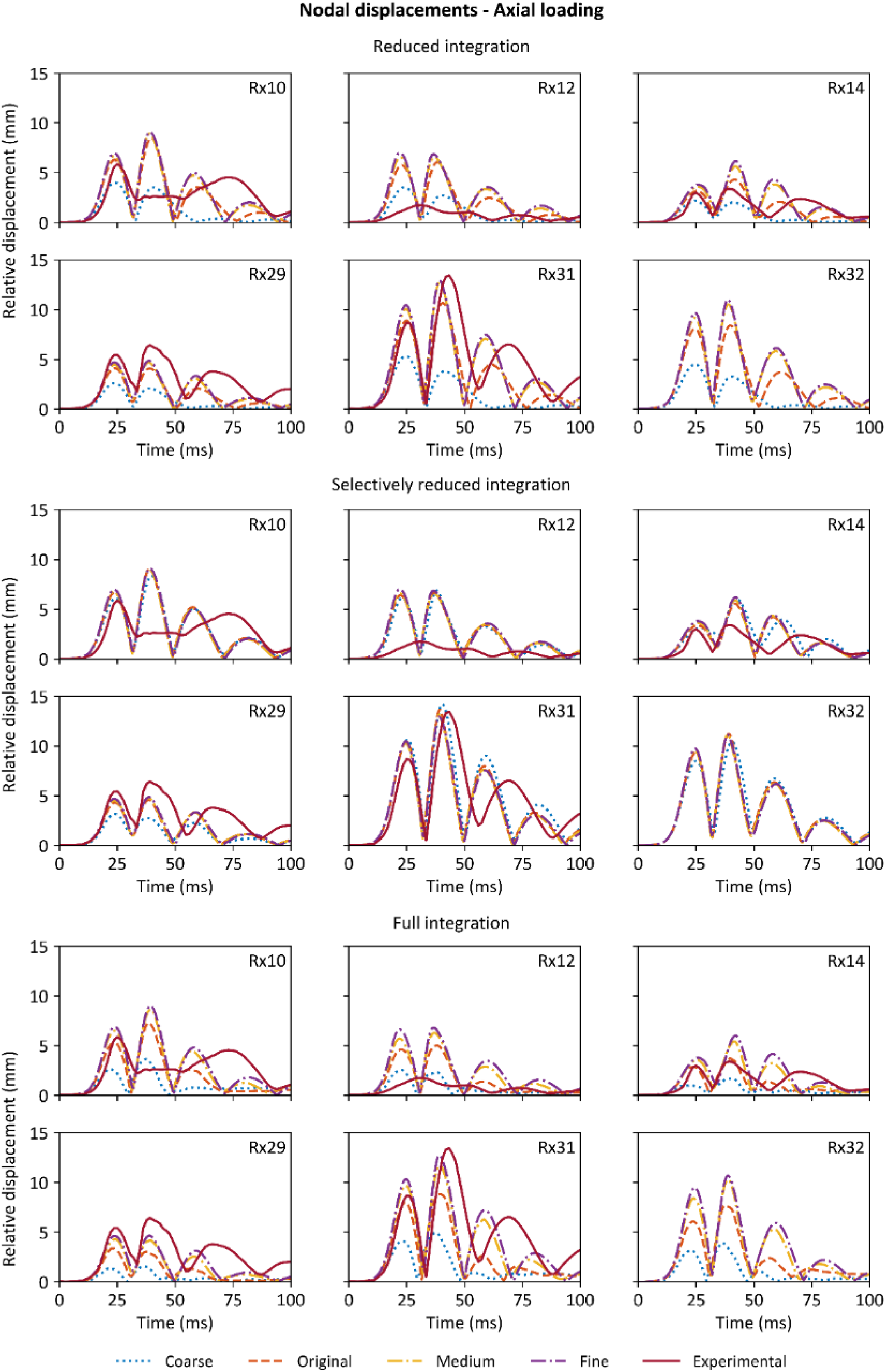
Time-history nodal displacements of the nodes of interest for the axial loading, with experimental results [43] for reference (no experimental results available for Rx32).

**Figure B2.**
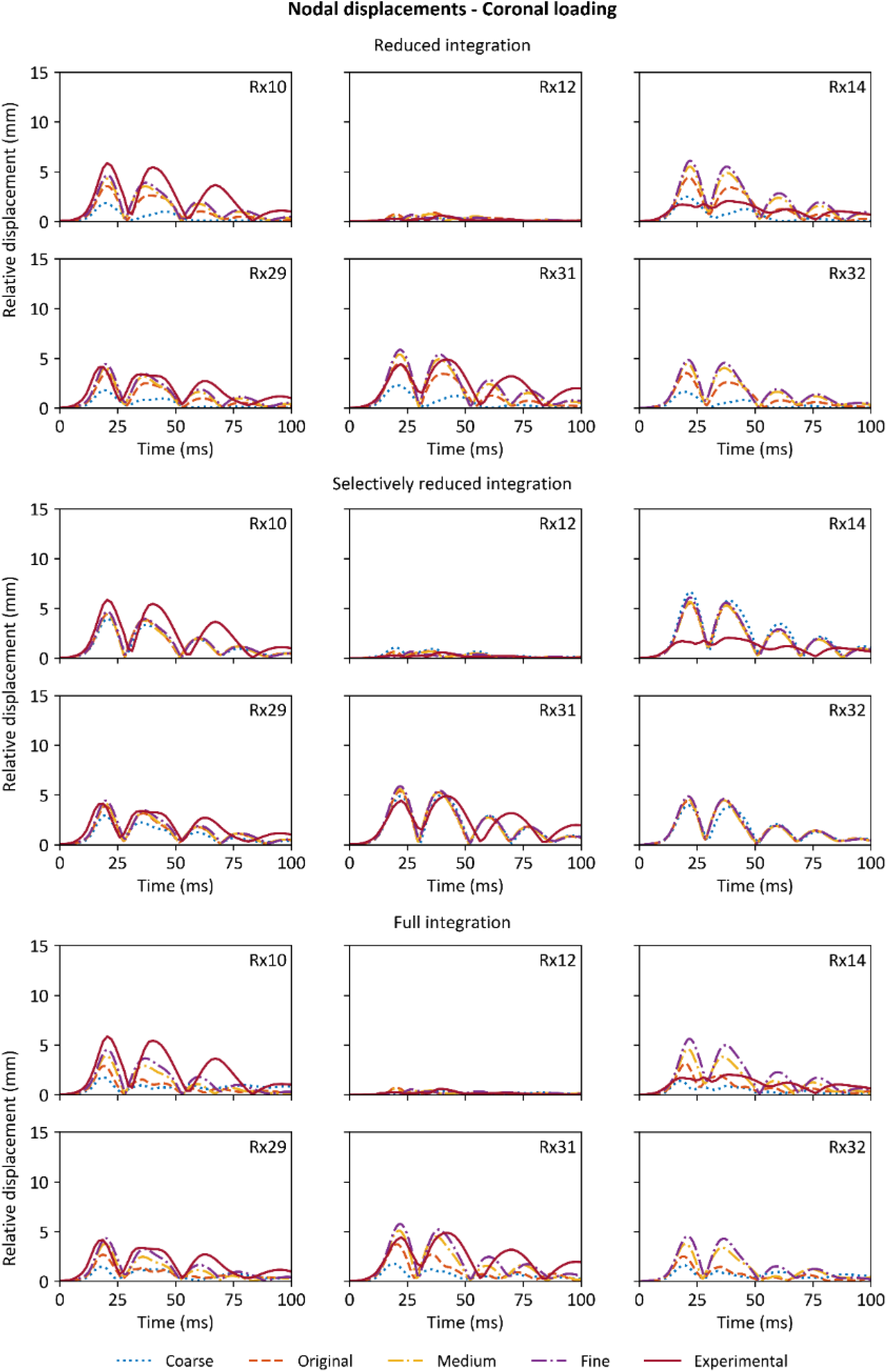
Time-history nodal displacements of the nodes of interest for the coronal loading, with experimental results [43] for reference (no experimental results available for Rx32).

**Figure B3.**
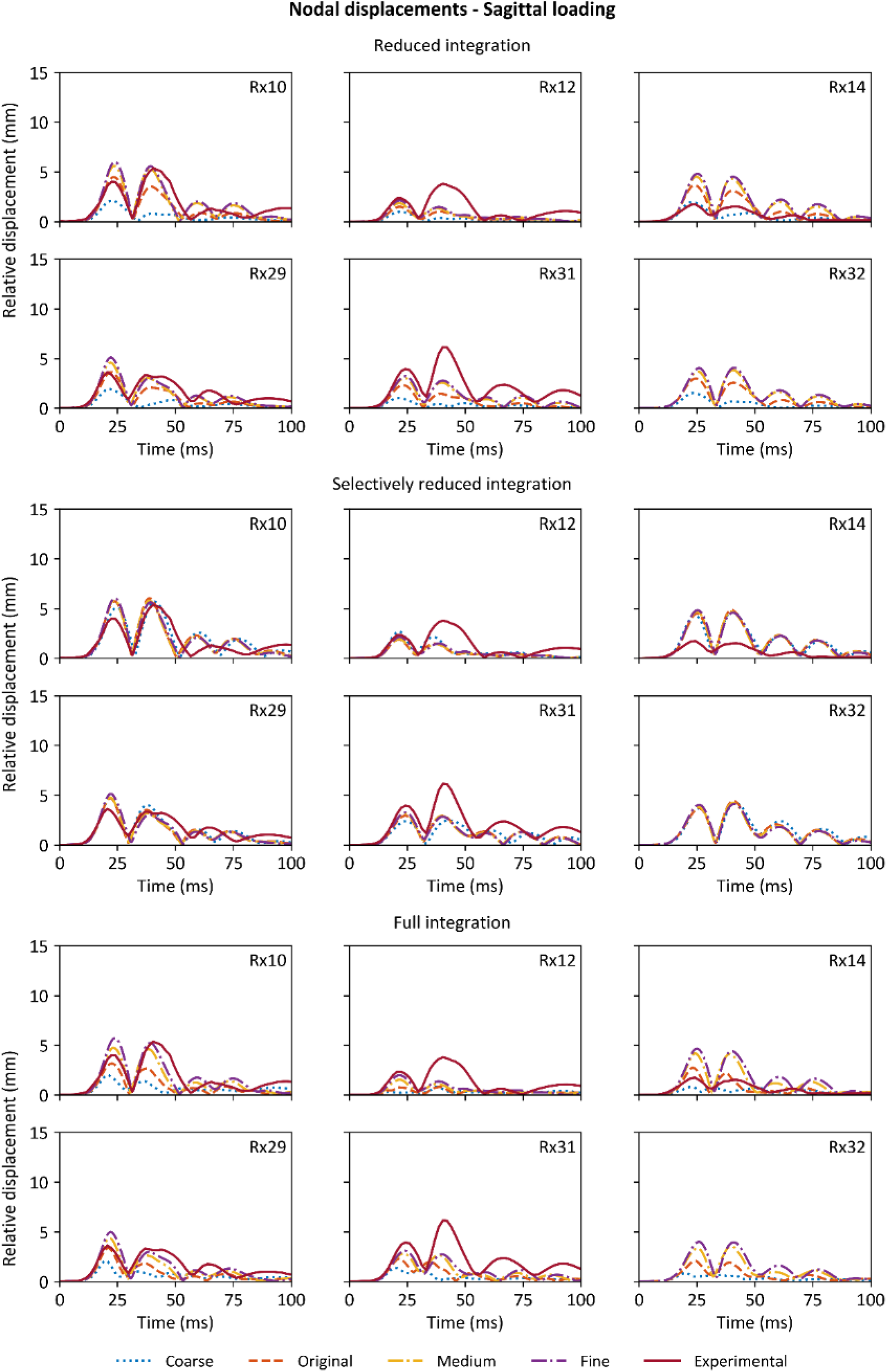
Time-history nodal displacements of the nodes of interest for the sagittal loading, with experimental results [43] for reference (no experimental results available for Rx32)

## Appendix C. Nodal displacement convergence tables

**Table C1.**
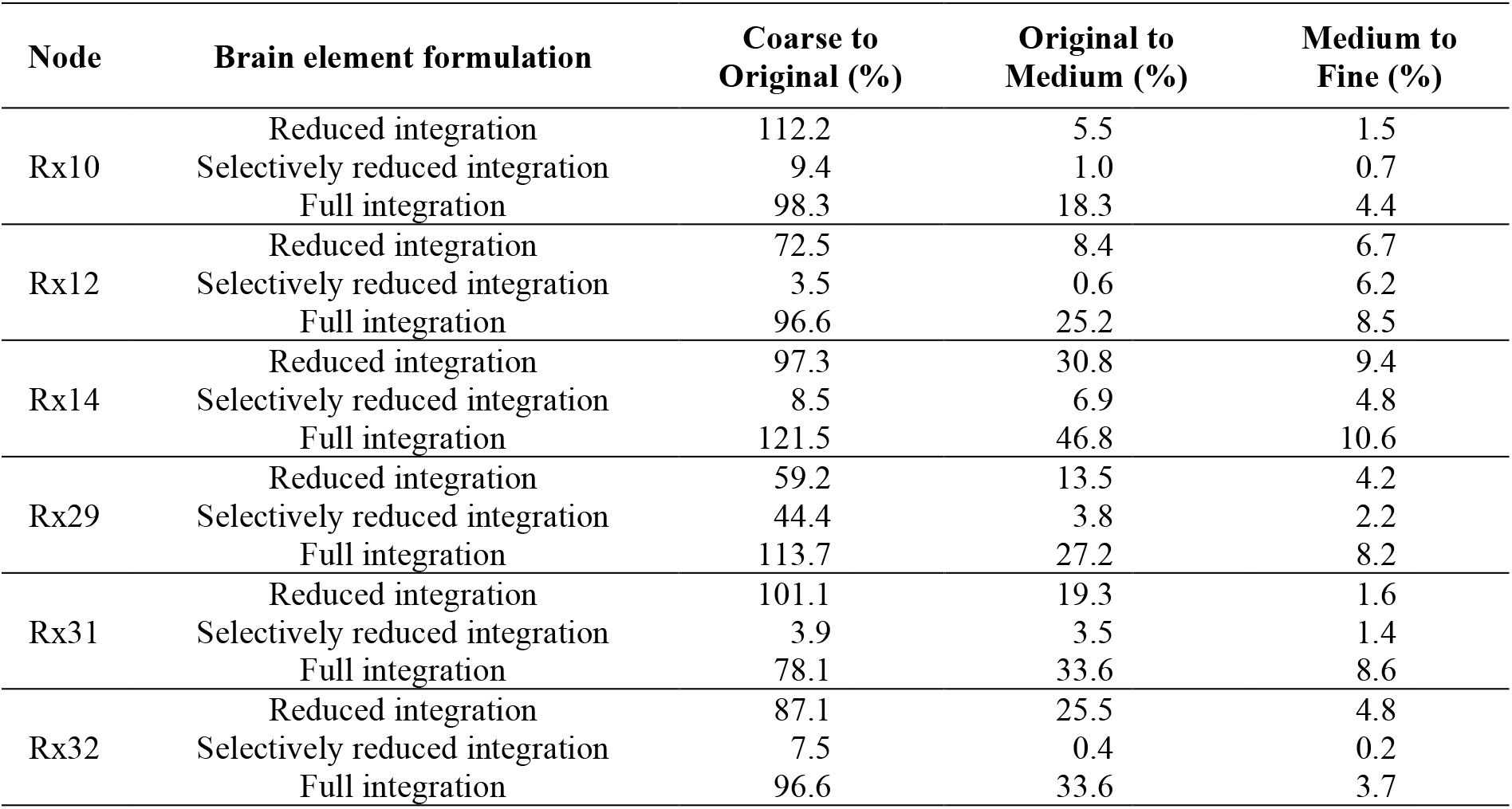
Absolute percentage of change for nodal displacements between two neighboring refinement steps, for the axial loading simulations.

**Table C2.** Absolute percentage of change for nodal displacements between two neighboring refinement steps, for the coronal loading simulations.

| Node | Brain element formulation | Coarse to Original (%) | Original to Medium (%) | Medium to Fine (%) |
| --- | --- | --- | --- | --- |
| Rx10 | Reduced integration | 95.7 | 21.0 | 8.1 |
|  | Selectively reduced integration | 11.6 | 0.8 | 5.3 |
|  | Full integration | 66.5 | 35.1 | 15.5 |
| Rx12 | Reduced integration | 74.3 | 16.8 | 11.9 |
|  | Selectively reduced integration | 29.5 | 13.7 | 2.3 |
|  | Full integration | 77.5 | 14.9 | 12.0 |
| Rx14 | Reduced integration | 76.4 | 24.9 | 11.0 |
|  | Selectively reduced integration | 17.4 | 3.0 | 7.8 |
|  | Full integration | 125.9 | 45.2 | 22.4 |
| Rx29 | Reduced integration | 92.9 | 17.9 | 9.0 |
|  | Selectively reduced integration | 32.5 | 6.0 | 6.0 |
|  | Full integration | 84.3 | 40.4 | 14.8 |
| Rx31 | Reduced integration | 94.5 | 21.1 | 8.8 |
|  | Selectively reduced integration | 6.0 | 3.6 | 5.7 |
|  | Full integration | 109.3 | 36.3 | 13.4 |
| Rx32 | Reduced integration | 113.4 | 25.7 | 9.1 |
|  | Selectively reduced integration | 13.1 | 2.9 | 5.5 |
|  | Full integration | 54.6 | 50.4 | 20.5 |

**Table C3.**
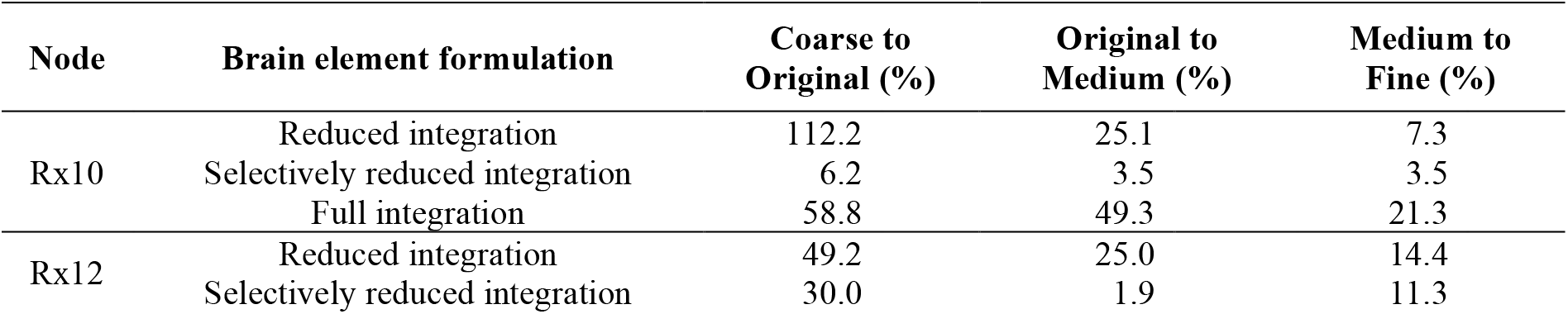
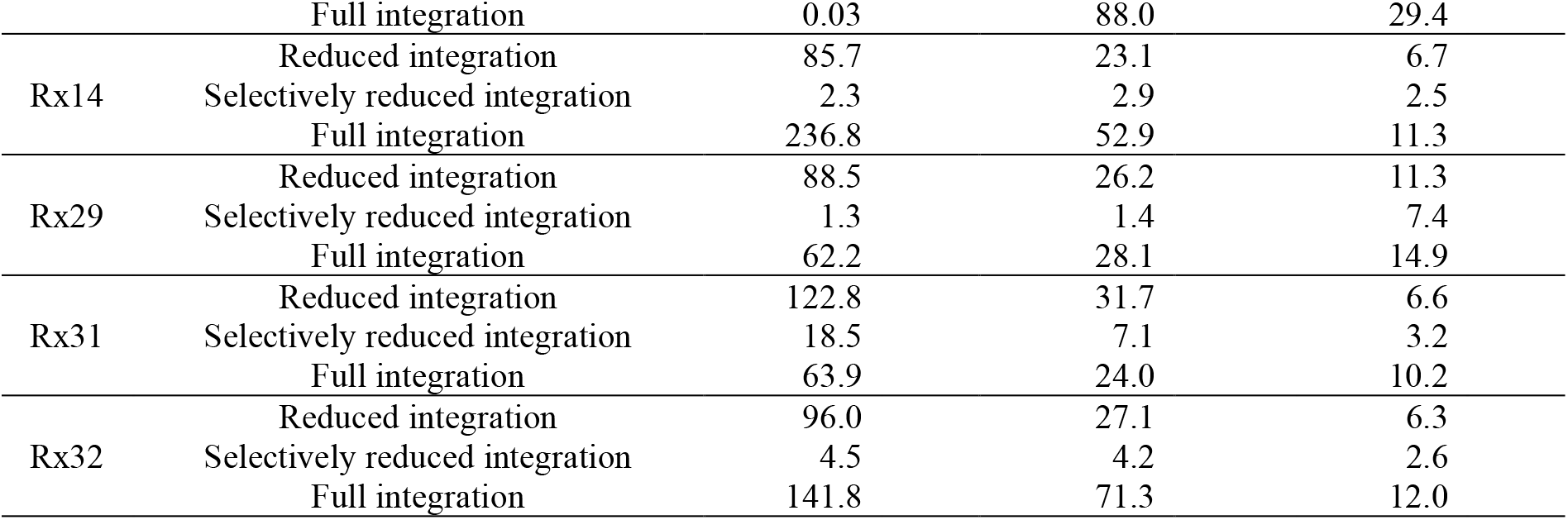
Absolute percentage of change for nodal displacements between two neighboring refinement steps, for the sagittal loading simulations.

## Appendix D. 95^th^ percentage maximum principal strain convergence table

**Table D1.**
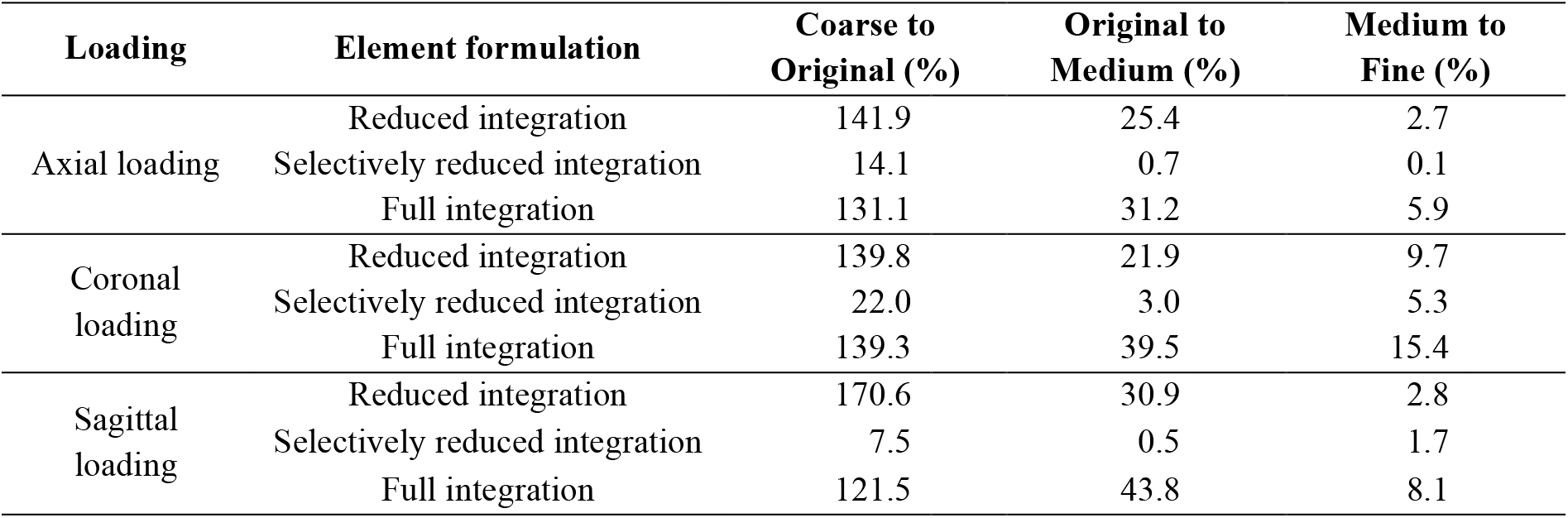
Absolute percentage of variation of the 95^th^ percentile maximum principal strain across the whole brain between two successive refinement steps of the brain mesh.

## Appendix E. Influence of the element formulation on the model convergence behavior based on the cumulative percentile strain

Figure E1 illustrates the influence of element formulation on the model convergence behavior based on the strain value of varying percentiles. For all three loading directions, the cumulative percentile strain curves for the different mesh densities were visually more separated with the reduced and full integration than with the S/R integration. When implementing the S/R integration, the cumulative percentile strain curves for the original, medium, and fine meshes were mostly overlapping, which reinforced that the S/R integration contributes to the fastest mesh convergence behavior than the reduced and full integration.

**Figure E1.**
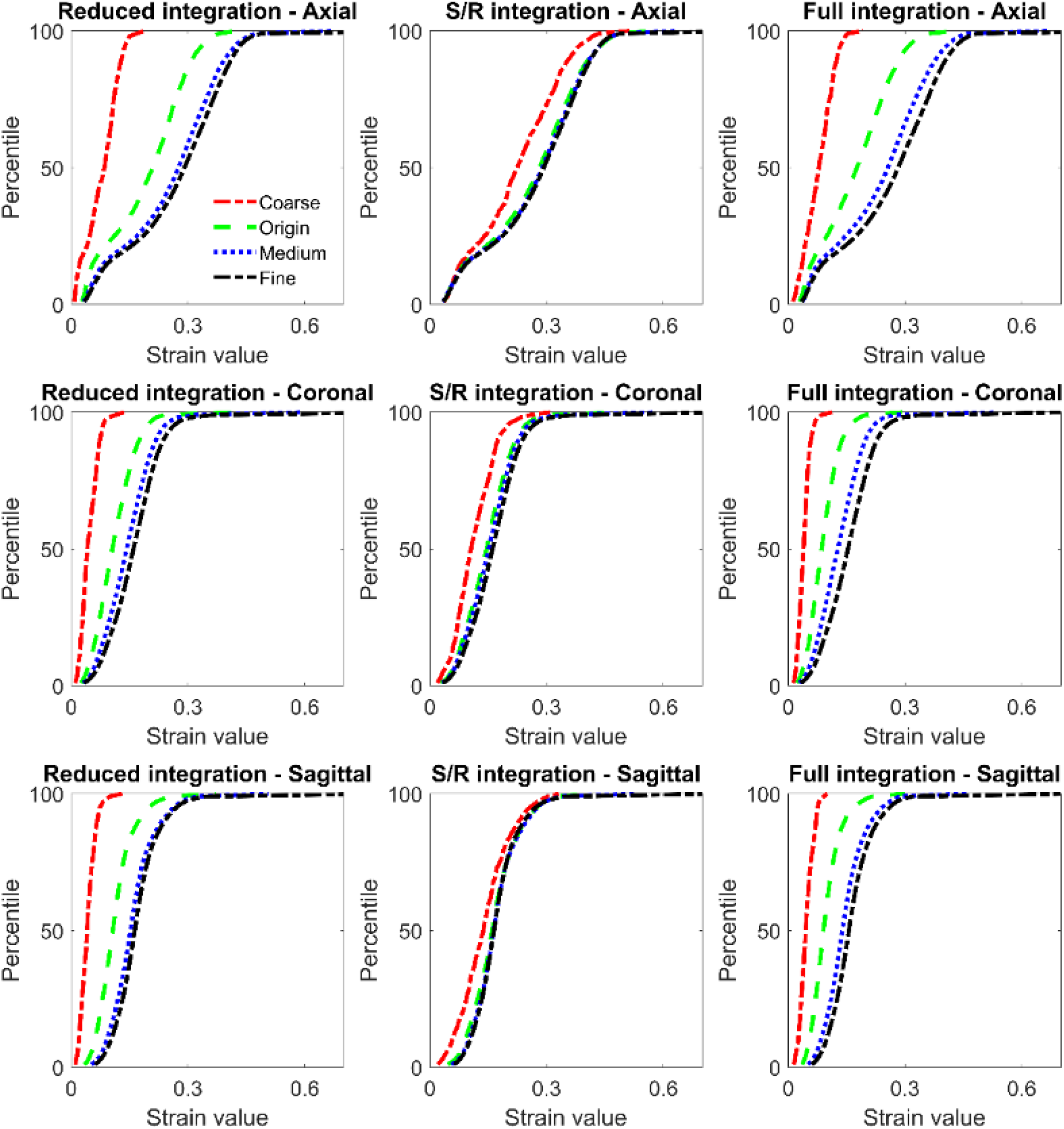
The cumulative percentile strain curves illustrate that the selectively reduced (S/R) integration (middle column) contributes to a fastest mesh convergence behavior than the reduced (left column) and full (right column) integration in all three loading conditions (i.e., axial, coronal and sagittal rotation in the top, middle, and low row, respectively).

## References

[1] M.C. Dewan, A. Rattani, S. Gupta, R.E. Baticulon, Y.-C. Hung, M. Punchak, A. Agrawal, A.O. Adeleye, M.G. Shrime, A.M. Rubiano, J.V. Rosenfeld, K.B. Park, Estimating the global incidence of traumatic brain injury, Journal of Neurosurgery, 130 (2018) 1080–1097.

[2] A.I.R. Maas, D.K. Menon, P.D. Adelson, N. Andelic, M.J. Bell, A. Belli, P. Bragge, A. Brazinova, A. Büki, R.M. Chesnut, G. Citerio, M. Coburn, D.J. Cooper, A.T. Crowder, E. Czeiter, M. Czosnyka, R. Diaz-Arrastia, J.P. Dreier, A.-C. Duhaime, A. Ercole, T.A. van Essen, V.L. Feigin, G. Gao, J. Giacino, L.E. Gonzalez-Lara, R.L. Gruen, D. Gupta, J.A. Hartings, S. Hill, J.-Y. Jiang, N. Ketharanathan, E.J.O. Kompanje, L. Lanyon, S. Laureys, F. Lecky, H. Levin, H.F. Lingsma, M. Maegele, M. Majdan, G. Manley, J. Marsteller, L. Mascia, C. McFadyen, S. Mondello, V. Newcombe, A. Palotie, P.M. Parizel, W. Peul, J. Piercy, S. Polinder, L. Puybasset, T.E. Rasmussen, R. Rossaint, P. Smielewski, J. Söderberg, S.J. Stanworth, M.B. Stein, N. von Steinbüchel, W. Stewart, E.W. Steyerberg, N. Stocchetti, A. Synnot, B. Te Ao, O. Tenovuo, A. Theadom, D. Tibboel, W. Videtta, K.K.W. Wang, W.H. Williams, L. Wilson, K. Yaffe, InTbir Participants and Investigators, Traumatic brain injury: integrated approaches to improve prevention, clinical care, and research, Lancet Neurol, 16 (2017) 987–1048.

[3] K. Jones, A. Theadom, N. Starkey, I. Zeng, S. Ameratunga, S. Barker-Collo, L. Wilkinson-Meyers, B.T. Ao, N. Henry, L.A. McClean, J. Chua, L. Haumaha, M. Kahan, G. Christey, N. Hardaker, A. Jones, A. Dowell, V. Feigin, K. Jones, A. Theadom, N. Starkey, S. Barker-Collo, M. Kahan, G. Christey, N. Hardaker, A. Jones, A. Dowell, V. Feigin, L. Wilkinson-Meyers, B.T. Ao, S. Ameratunga, I. Zeng, J. Chua, L. Haumaha, N. Henry, L.A. McClean, K. Berryman, N. Scott, B. Masters-Awatere, A population-based study of traumatic brain injury incidence and mechanisms in New Zealand: 2021–2022 compared with 2010–2011, The Lancet Regional Health – Western Pacific, 67 (2026).

[4] J.-C. Xu, Z. Zhou, C. Ahlström, S. Kleiven, Incidence of Traumatic Head Injuries in Swedish Specialised Care from 2008-2022, in: International Research Council on the Biomechanics of Injury, Vilnius, Lithuania, 2025.

[5] S. Meng, R. Schindler, S. Kleiven, N. Lubbe, Traumatic Brain Injury in Vulnerable Road Users: Analysis of German In-Depth Crash Data to Inform Targeted Prevention, Stapp Car Crash Journal, 70 (2026) 42–53.

[6] J. Ho, S. Kleiven, Can sulci protect the brain from traumatic injury?, Journal of Biomechanics, 42 (2009) 2074–2080.

[7] C. Li, Z. Zhou, The influence of tension-compression switches on brain anisotropic modelling, International Journal of Engineering Science, 227 (2026) 104596.

[8] Z. Zhou, X. Li, S. Kleiven, Fluid–structure interaction simulation of the brain–skull interface for acute subdural haematoma prediction: Z. Zhou et al, Biomech Model Mechanobiol, 18 (2019) 155–173.

[9] Z. Zhou, X. Li, S. Kleiven, C.S. Shah, W.N. Hardy, A reanalysis of experimental brain strain data: implication for finite element head model validation, SAE Technical Paper, (2018).

[10] S. Ji, M. Ghajari, H. Mao, R.H. Kraft, M. Hajiaghamemar, M.B. Panzer, R. Willinger, M.D. Gilchrist, S. Kleiven, J.D. Stitzel, Use of brain biomechanical models for monitoring impact exposure in contact sports, Ann Biomed Eng, 50 (2022) 1389–1408.

[11] P.V. Bayly, E.H. Clayton, G.M. Genin, Quantitative imaging methods for the development and validation of brain biomechanics models, Annual review of biomedical engineering, 14 (2012) 369–396.

[12] S. Kleiven, W.N. Hardy, Correlation of an FE Model of the Human Head with Local Brain Motion--Consequences for Injury Prediction, Stapp Car Crash Journal, 46 (2002) 123–144.

[13] W. Zhao, S. Ji, Mesh Convergence Behavior and the Effect of Element Integration of a Human Head Injury Model, Ann Biomed Eng, 47 (2019) 475–486.

[14] S. Kleiven, H. von Holst, Consequences of head size following trauma to the human head, Journal of Biomechanics, 35 (2002) 153–160.

[15] H. Mao, H. Gao, L. Cao, V.V. Genthikatti, K.H. Yang, Development of high-quality hexahedral human brain meshes using feature-based multi-block approach, Computer Methods in Biomechanics and Biomedical Engineering, 16 (2013) 271–279.

[16] E.G. Takhounts, R.H. Eppinger, J.Q. Campbell, R.E. Tannous, E.D. Power, L.S. Shook, On the Development of the SIMon Finite Element Head Model, Stapp Car Crash Journal, 47 (2003) 107–133.

[17] J.S. Giudice, W. Zeng, T. Wu, A. Alshareef, D.F. Shedd, M.B. Panzer, An Analytical Review of the Numerical Methods used for Finite Element Modeling of Traumatic Brain Injury, Ann Biomed Eng, 47 (2019) 1855–1872.

[18] H.T. Garimella, R.H. Kraft, Modeling the mechanics of axonal fiber tracts using the embedded finite element method, International Journal for Numerical Methods in Biomedical Engineering, 33 (2017) e2823.

[19] R.D. Cook, D.S. Malkus, M.E. Plesha, Concepts and applications of finite element analysis, 3. ed., Wiley, New York, 1989.

[20] D.S. Bombarde, L.N. Silla, S.S. Gautam, A. Nandy, A Comprehensive Comparative Review of Various Advanced Finite Elements to Alleviate Shear, Membrane and Volumetric Locking, Arch Computat Methods Eng, 31 (2024) 1979–2013.

[21] S. Kleiven, Predictors for traumatic brain injuries evaluated through accident reconstructions, Stapp Car Crash Journal, 51 (2007) 81–114.

[22] R.R.L. Hosey, Y. K., A homeomorphic finite element model of the human head and neck, in: B.R.G. Simon, R. H.; Johnson, P. C.; Gross, J. F. (Ed.) Finite Elements in Biomechanics, Wiley & Son, 1981, pp. 379–401.

[23] C.C. Ward, R.B. Thompson, The Development of a Detailed Finite Element Brain Model, SAE Transactions, 84 (1975) 3238–3252.

[24] D. Sahoo, C. Deck, R. Willinger, Development and validation of an advanced anisotropic visco-hyperelastic human brain FE model, Journal of the Mechanical Behavior of Biomedical Materials, 33 (2014) 24–42.

[25] H. Mao, L. Zhang, B. Jiang, V.V. Genthikatti, X. Jin, F. Zhu, R. Makwana, A. Gill, G. Jandir, A. Singh, K.H. Yang, Development of a Finite Element Human Head Model Partially Validated With Thirty Five Experimental Cases, J Biomech Eng, 135 (2013).

[26] L. Zhang, K.H. Yang, A.I. King, Comparison of Brain Responses Between Frontal and Lateral Impacts by Finite Element Modeling, Journal of Neurotrauma, 18 (2001) 21–30.

[27] E.G. Takhounts, S.A. Ridella, V. Hasija, R.E. Tannous, J.Q. Campbell, D. Malone, K. Danelson, J. Stitzel, S. Rowson, S. Duma, Investigation of Traumatic Brain Injuries Using the Next Generation of Simulated Injury Monitor (SIMon) Finite Element Head Model, Stapp Car Crash Journal, 52 (2008) 1–31.

[28] S. Ji, W. Zhao, J.C. Ford, J.G. Beckwith, R.P. Bolander, R.M. Greenwald, L.A. Flashman, K.D. Paulsen, T.W. McAllister, Group-Wise Evaluation and Comparison of White Matter Fiber Strain and Maximum Principal Strain in Sports-Related Concussion, Journal of Neurotrauma, 32 (2015) 441–454.

[29] H. Kimpara, Y. Nakahira, M. Iwamoto, K. Miki, K. Ichihara, S.-i. Kawano, T. Taguchi, Investigation of Anteroposterior Head-Neck Responses during Severe Frontal Impacts Using a Brain-Spinal Cord Complex FE Model, Stapp Car Crash Journal, 50 (2006) 509–544.

[30] H. Duckworth, A. Azor, N. Wischmann, K.A. Zimmerman, I. Tanini, D.J. Sharp, M. Ghajari, A Finite Element Model of Cerebral Vascular Injury for Predicting Microbleeds Location, Front. Bioeng. Biotechnol., 10 (2022).

[31] A. Alshareef, A.K. Knutsen, C.L. Johnson, A. Carass, K. Upadhyay, P.V. Bayly, D.L. Pham, J.L. Prince, K.T. Ramesh, Integrating material properties from magnetic resonance elastography into subject-specific computational models for the human brain, Brain Multiphysics, 2 (2021) 100038.

[32] E. Griffiths, J. Hinrichsen, N. Reiter, S. Budday, On the importance of using region-dependent material parameters for full-scale human brain simulations, European Journal of Mechanics - A/Solids, 99 (2023) 104910.

[33] J. Ho, H. von Holst, S. Kleiven, Automatic generation and validation of patient-specific finite element head models suitable for crashworthiness analysis, International Journal of Crashworthiness, 14 (2009) 555–563.

[34] X. Li, Z. Zhou, S. Kleiven, An anatomically detailed and personalizable head injury model: Significance of brain and white matter tract morphological variability on strain, Biomech Model Mechanobiol, 20 (2021) 403–431.

[35] Z. Zhou, X. Li, S. Kleiven, Biomechanics of periventricular injury, Journal of Neurotrauma, 37 (2020) 1074–1090.

[36] Z. Zhou, X. Li, A.G. Domel, E.L. Dennis, M. Georgiadis, Y. Liu, S.J. Raymond, G. Grant, S. Kleiven, D. Camarillo, The presence of the temporal horn exacerbates the vulnerability of hippocampus during head impacts, Front. Bioeng. Biotechnol., 10 (2022) 754344.

[37] Ansys, LS-DYNA Theory Manual, R15.0, Ansys, Inc., 2024.

[38] D.P. Flanagan, T. Belytschko, A uniform strain hexahedron and quadrilateral with orthogonal hourglass control, International Journal for Numerical Methods in Engineering, 17 (1981) 679–706.

[39] Z. Zhou, X. Li, S. Kleiven, Surface-based versus voxel-based finite element head models: comparative analyses of strain responses, Biomech Model Mechanobiol, 24 (2025) 845–864.

[40] J.O. Hallquist, LS-DYNA Theory Manual, Livermore Software Technology Corp (LSTC), 2006.

[41] R. van Noort, M.M. Black, T.R.P. Martin, S. Meanley, A study of the uniaxial mechanical properties of human dura mater preserved in glycerol, Biomaterials, 2 (1981) 41–45.

[42] P. Aimedieu, R. Grebe, Tensile strength of cranial pia mater: preliminary results, Journal of Neurosurgery, 100 (2004) 111–114.

[43] A. Alshareef, J.S. Giudice, J. Forman, D.F. Shedd, K.A. Reynier, T. Wu, S. Sochor, M.R. Sochor, R.S. Salzar, M.B. Panzer, Biomechanics of the Human Brain during Dynamic Rotation of the Head, Journal of Neurotrauma, 37 (2020) 1546–1555.

[44] M.B. Panzer, B.S. Myers, B.P. Capehart, C.R. Bass, Development of a Finite Element Model for Blast Brain Injury and the Effects of CSF Cavitation, Ann Biomed Eng, 40 (2012) 1530–1544.

[45] S. Sullivan, S.A. Eucker, D. Gabrieli, C. Bradfield, B. Coats, M.R. Maltese, J. Lee, C. Smith, S.S. Margulies, White matter tract-oriented deformation predicts traumatic axonal brain injury and reveals rotational direction-specific vulnerabilities, Biomech Model Mechanobiol, 14 (2015) 877–896.

[46] T. Wu, A. Alshareef, J.S. Giudice, M.B. Panzer, Explicit Modeling of White Matter Axonal Fiber Tracts in a Finite Element Brain Model, Ann Biomed Eng, 47 (2019) 1908–1922.

[47] O.C. Zienkiewicz, R.L. Taylor, The Finite Element Method: Its Basis and Fundamentals, 5th ed., 2000.

[48] D.S. Malkus, T.J.R. Hughes, Mixed finite element methods — Reduced and selective integration techniques: A unification of concepts, Computer Methods in Applied Mechanics and Engineering, 15 (1978) 63–81.

[49] T.J.R. Hughes, M. Cohen, M. Haroun, Reduced and selective integration techniques in the finite element analysis of plates, Nuclear Engineering and Design, 46 (1978) 203–222.

[50] A. Alshareef, A. Carass, Y.-C. Lu, J. Mojumder, A.M. Diano, O.M. Bailey, R.J. Okamoto, D.L. Pham, J.L. Prince, P.V. Bayly, C.L. Johnson, Average Biomechanical Responses of the Human Brain Grouped by Age and Sex, Ann Biomed Eng, 53 (2025) 1496–1511.

[51] Z. Zhou, C. Olsson, T.C. Gasser, X. Li, S. Kleiven, The White Matter Fiber Tract Deforms Most in the Perpendicular Direction During *In Vivo* Volunteer Impacts, Journal of Neurotrauma, 41 (2024) 2554–2570.

